# Quantitative PCR assays for detection of five Arctic fish species: *Lota lota, Cottus cognatus, Salvelinus alpinus, Salvelinus malma*, and *Thymallus arcticus* from environmental DNA

**DOI:** 10.1101/116053

**Authors:** Torrey W. Rodgers, John. R. Olson, Stephen L. Klobucar, Karen. E. Mock

## Abstract

The North Slope of Alaska contains arctic fish populations that are important for subsistence of local human populations, and are under threat from natural resource extraction and climate change. We designed and evaluated four quantitative PCR assays for the detection of environmental DNA from five Alaskan fish species present on the North Slope of Alaska: burbot (*Lota lota*), arctic char (*Salvelinus alpinus*), Dolly Varden (*Salvelinus malma*), arctic grayling (*Thymallus arcticus*), and slimy sculpin (*Cottus cognatus*). All assays were designed and tested for species specificity and sensitivity, and all assays detected target species from filtered water samples collected from the field. These assays will enable efficient and economical detection and monitoring of these species in lakes and rivers. This in turn will provide managers with improved knowledge of current distributions and future range shifts associated with climate and development threats, enabling more timely management.

## Introduction

The Arctic is warming faster than any other region on the globe (Reist et al. 2006) with a 2.1 °C increase in air temperature observed over the last 30 years (ACIA 2005). On the North Slope of Alaska (hereafter the North Slope), a vast region encompassing over 245,000 km^2^ including the Arctic National Wildlife Refuge, Prudhoe Bay, and the National Petroleum Reserve-Alaska, climate change has, and will continue to have, a disproportionally large and rapid impact compared to the continental U.S (Stewart et al. 2013). Moreover, across the North Slope, native fish species are an important source of subsistence for local human populations including Inupiat and Nunamiut communities (Courtney et al. 2016; Moerlein and Carothers 2012). If aquatic food webs do not respond proportionally to the warming climate, local extinctions of fish populations could occur (Budy and Luecke 2014). Additionally, many North Slope fish populations have high potential for negative effects from natural resource extraction, as the North Slope holds a large portion of North America’s petroleum reserves (Council 2003). Thus, arctic fish and the local human communities that depend on them are exceedingly vulnerable (Himes-Cornell and Kasperski 2015).

Given the ecological risks and array of conflicting human interests in this region, accurate monitoring and assessment of native North Slope fish populations is critical for effective management and policy decisions. Accurate assessment and monitoring of these fish species, however, is logistically challenging and expensive. In some areas, freshwater covers as much as 48% of the land surface (Riordan et al. 2006), sampling is restricted to summer months, and sites are only accessible by air. Conducting surveys using traditional techniques such as gill or trap nets requires transportation of bulky gear as well as extensive man hours, severely limiting the number of sites visited in a season. Thus, there is a need for accurate techniques capable of rapidly detecting fish species of concern in order to increase the number of sites that can be sampled, and the amount of meaningful data that can be collected for these species.

Environmental DNA (eDNA) detection is a molecular technique that uses trace amounts of DNA from the water column to detect aquatic species including fish (Jerde et al. 2011; Takahara et al. 2013). eDNA has been used for the detection of many fish species worldwide, and has been shown to be more sensitive than traditional sampling (Janosik and Johnston 2015; Wilcox et al. 2016). In addition, an eDNA sample can be collected by a single person in less than 30 minutes, greatly increasing the number of sites that can be sampled in a single field season. Thus, this technique is ideal for remote sites such as the North Slope of Alaska where sampling is logistically difficult and expensive.

We developed and tested four highly sensitive eDNA assays to detect five fish species native to the North Slope of Alaska: burbot (*Lota lota),* slimy sculpin *(Cottus cognatus),* arctic char *(Salvelinus alpinus),* Dolly Varden *(Salvelinus malma),* and arctic grayling *(Thymallus arcticus)*. These assays use Taqman^®^-based quantitative PCR (qPCR) to detect species from filtered water samples. Use of these assays will improve knowledge of species distributions on the North Slope as well as throughout Alaska and the Arctic. In addition, the *L. lota* assay may be useful for early detection of this species outside of Alaska where it is invasive (Gardunio et al. 2011). These assays will allow for improved monitoring of distributional shifts, and generation of presence/absence data can be used to inform models that predict species responses to natural resource extraction and climate change.

## Methods

### Primer and probe development

Reference sequences from the mitochondrial genes cytochrome oxidase subunit 1 (COI) and cytochrome b (cytb) for *L. lota, C. cognatus, S. alpinus, S. malma, T. arcticus* and 22 other potentially sympatric fish species from Alaska (Supplementary Table S1) were obtained by Sanger sequencing using primers and conditions from Crete-Lafreniere et al. (2012), with the exception of cytb for *C. cognatus* which was sequenced using the custom designed primers Cco-F 5’-GCC AGC CTA CGA AAA ACC CA- 3’ and Cco-R 5’- TCT ATT CAG CCT GCT ATT GGG A -3’. Reference samples were obtained from the University of Alaska museum (UAM), the Bureau of Land Management (BLM) Alaska field office, the United State Fish and Wildlife Service (USFWS), and the Utah State University (USU) Fish Ecology Lab. Sequences have been deposited into NCBI Genbank with accession numbers in Table S1. In addition, for *C. cognatus* assay design we included reference sequences from Genbank for two *Cottus* species present in coastal Alaska (but not on the North Slope) for which we could not obtain tissue samples: *C. aleuticus* (accession # AF549106), and *C. asper* (accession # AF549105).

Sequences from all species were aligned using Sequencher software (Gene Codes; Ann Arbor, MI). We used the online tool DECEPHIR (Wright et al. 2014) to select species specific primers that would amplify target species while excluding all sympatric non-target species. For *S. alpinus,* and *S. malma*, no primer sites could be identified with enough polymorphism for species specificity due to the fact that these two species have extremely little mitochondrial variation (justification for species delineation is based on morphology, ecology, and nuclear DNA (Taylor et al. 2008)). Thus, we designed one primer set that can detect both *S. alpinus and S. malma* but cannot distinguish between them, while still excluding all other species. For the remaining species, assays were designed to be species-specific. We used ABI primer express software (applied Biosystems; Foster City, CA) to design Taqman^®^ Minor Groove Binding qPCR probes with at least one mismatch to all non-target species, and also to modify primer length to meet melting temperature requirements for Taqman^®^ qPCR where necessary. This resulted in an assay for *L. lota* that amplifies 91 base pairs of cytb, an assay for *C. cognatus* that amplifies 118 base pairs of cytb, an assay for *S. alpinus/S. malma* that amplifies 145 base pairs of cytb, and an assay for *T. arcticus* that amplifies 116 base pairs of COI. Primer and probe sequences are presented in Table 1.

**Table 1.**
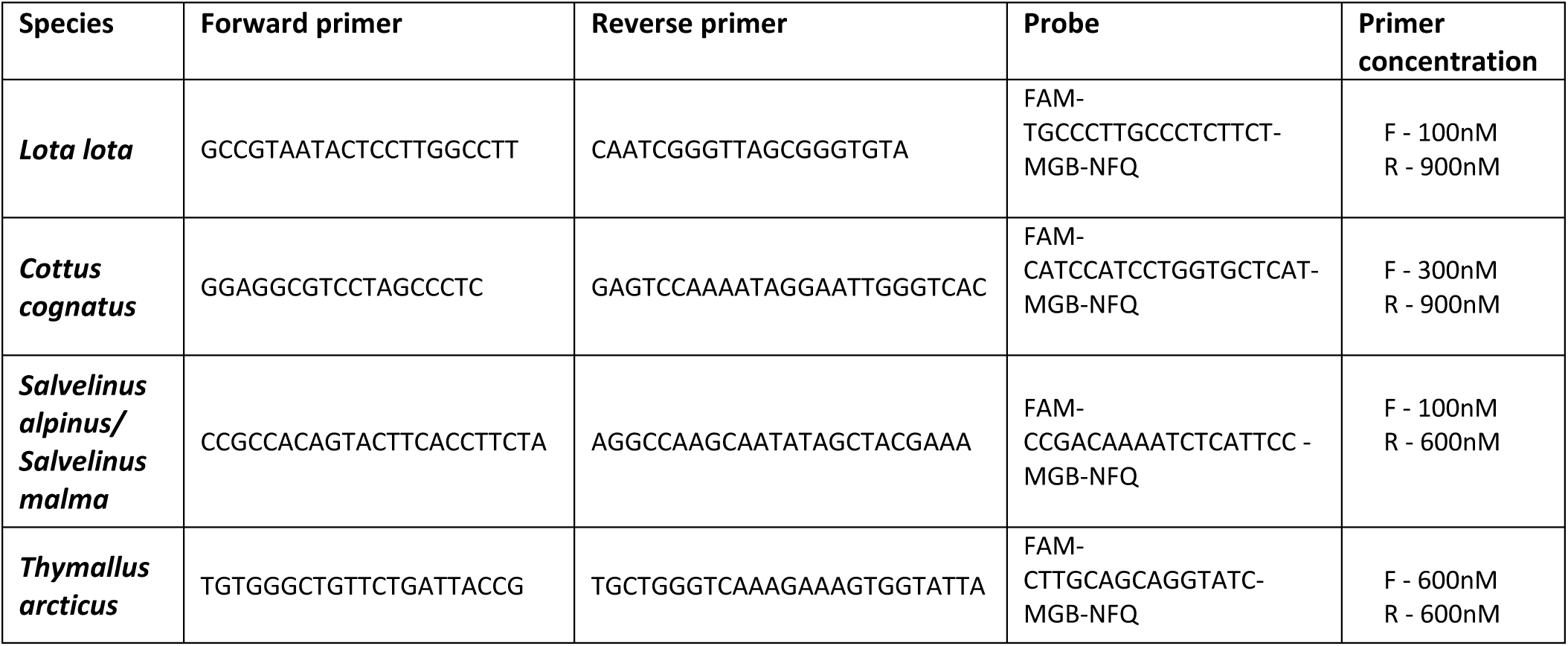
Primer and probe sequences and optimized primer concentrations for environmental DNA detection of 5 Alaskan fish species by taqman qPCR. All sequences are 5’ to 3’.

### Assay specificity testing

Primer sets were first tested for target amplification, species specificity, and primer dimer formation with SYBR^®^-Green qPCR. Primer sets were tested on all corresponding target species samples from Table S1, as well as on closely related non-target species samples (for salmonids, all other salmonids, for non-salmonids, all other non-salmonids from Table S1). qPCR reactions included 10 μl Power SYBR^®^-green Master Mix (Thermo-Fisher; Waltham, MA) 900nM of each primer, 4.4 μl of sterile H_2_O and 0.2ng of template DNA in a total reaction volume of 20 μl. Cycling conditions were 95° C for 10 minutes followed by 45 cycles of 95° C for 15 seconds and 60° C for one minute, followed by a melting curve to test for primer dimers and off target amplification.

Next, all primer sets and probes were tested in Taqman^®^ qPCR. Testing included all corresponding target species samples, at least one sample for each non-target species from table S1, and all closely related non-target species samples from Table S1 (for salmonid assays, all other salmonids, for non-salmonid assays, all other non-salmonids). qPCR reaction included 10 μl Taqman^®^ Environmental Master Mix (Thermo-Fisher; Waltham, MA), 900nM of each primer, 250nM of probe, 7 μl of sterile H_2_O and 0.2ng of template DNA in a total reaction volume of 20 μl. Cycling conditions were 95° C for 10 minutes followed by 45 cycles of 95° C for 15 seconds and 60° C for one minute. Finally, primer concentrations were optimized by running permutations of forward and reverse primers at 100 nM, 300 nM, 600 nM and 900 nM, each in triplicate, with 0.1 ng of target species DNA per reaction. Primer concentrations with the greatest peak fluorescence and lowest C_t_ value were selected for further work (Table 1). Negative controls were included in all qPCR reactions to test for contamination at all stages.

Finally, to test for any potential cross-amplification from species not tested empirically, we conducted a Genbank search of species that could potentially cross-amplify with our primer sets using NCBI Primer-BLAST (Ye et al. 2012). We used the default settings: ‘primer must have at least 2 total mismatches to unintended targets, including at least 2 mismatches within the last 5 bps. at the 3' end, Ignore targets that have 6 or more mismatches to the primer’.

### Assay sensitivity testing

To test assay sensitivity, we ordered MiniGene plasmids for each assay from Integrated DNA Technologies (Coralville, Iowa, USA) containing the assay sequence. The plasmid was suspended in 100 μL of TE (10 mM Tris, 0.1 mM EDTA) buffer, linearized by digestion with the enzyme Pvu1, and purified with a PureLink PCR Micro Kit (Thermo-Fisher; Waltham, MA) following manufacturer protocols. The resulting product was quantified with a Qubit fluorometer, and concentration was converted to copy number based on molecular weight (Wilcox et al. 2013). For sensitivity testing the product was diluted to create quantities of 5, 10, 20, 50, and 100 copies/reaction. Each of these quantities was run in qPCR in 6 replicates to determine assay sensitivity. We also ran a 5X, 5 fold plasmid standard curve (10, 50, 250, 1250 and 6250 copies in triplicate) for each assay to determine amplification efficiency.

### Field Validation

All Taqman assays were tested on eDNA samples collected by filtration from the Alaskan North Slope where the target species were known to be present based on traditional surveys. Samples were collected from five known positive locations for *L. lota,* and 20 known positive locations for *T. arcticus* from the National Petroleum Reserve-Alaska, and *5* known positive locations for C. *cognatus and S. alpinus* from near the Toolik Field Station (Figure 1 and Table 2). In addition, we tested each assay on a minimum of 3 samples collected from locations where each species was known to be absent (outside of the species range).

**Figure 1.**
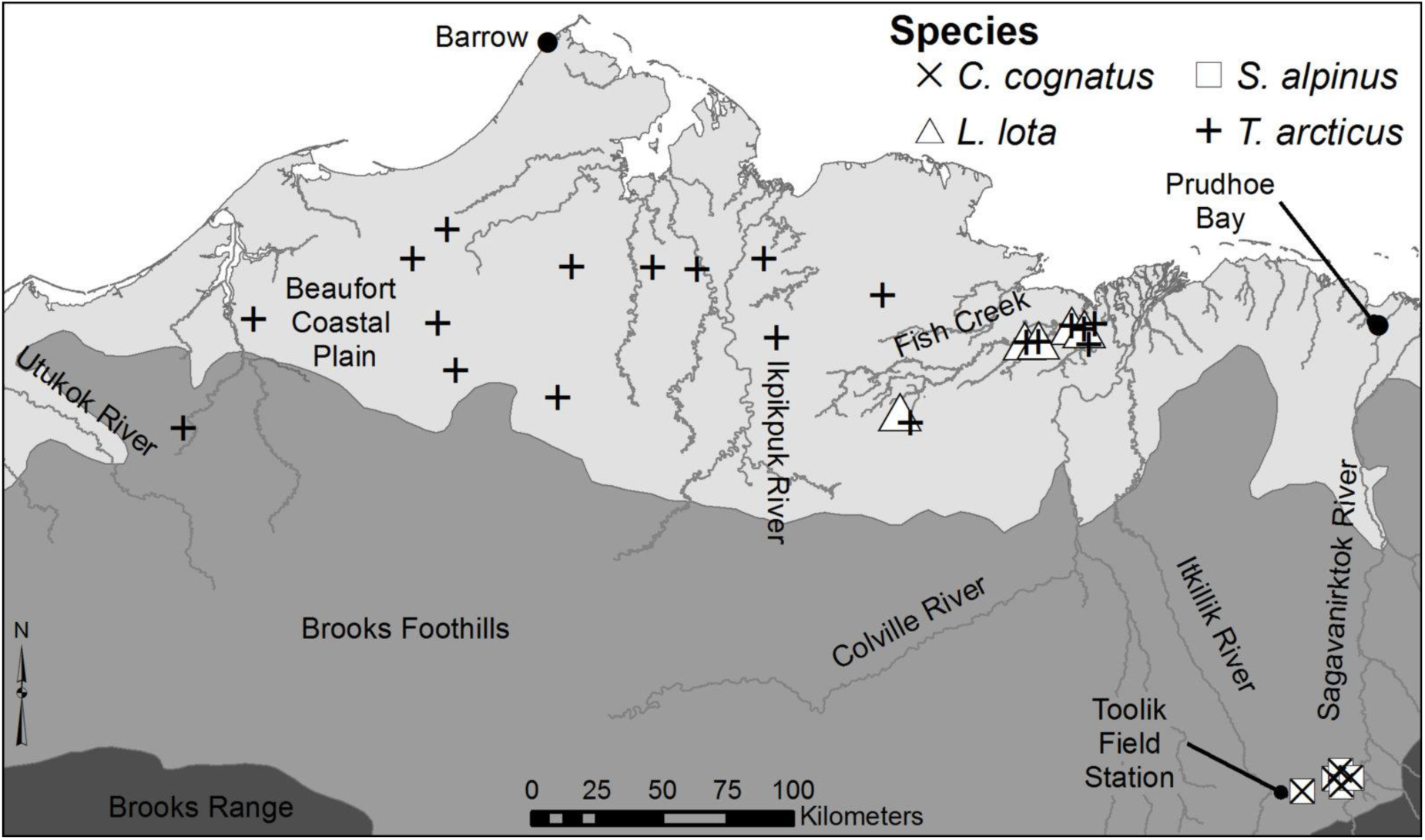
Map of locations in the North Slope of Alaska where eDNA samples were collected for field testing.

**Table 2.** Locations of eDNA samples collected for field validation of eDNA assays in the National Petroleum Reserve Alaska (NPRA) and Toolik Field Station (TFS). At each location, the species tested for with eDNA was known to be present based on traditional sampling.

| Species | Location | Collection date | Latitude | longitude | # of reactions amplified |
| --- | --- | --- | --- | --- | --- |
| *L. lota* | Unnamed creek, NPRA | 7/7/15 | 70.016 | -153.168 | 2/3 |
| *L. lota* | Unnamed creek, NPRA | 7/10/15 | 70.302 | -151.439 | 1/3 |
| *L. lota* | Unnamed creek, NPRA | 7/11/15 | 70.289 | -151.313 | 1/3 |
| *L. lota* | Unnamed creek, NPRA | 7/14/15 | 70.251 | -151.778 | 1/3 |
| *L. lota* | Unnamed creek, NPRA | 7/14/15 | 70.255 | -151.773 | 3/3 |
| *T. arcticus* | Unnamed creek, NPRA | 7/7/15 | 69.979 | -153.065 | 3/3 |
| *T. arcticus* | Unnamed creek, NPRA | 7/10/15 | 70.302 | -151.439 | 3/3 |
| *T. arcticus* | Unnamed creek, NPRA | 7/11/15 | 70.289 | -151.313 | 2/3 |
| *T. arcticus* | Unnamed creek, NPRA | 7/11/15 | 70.302 | -151.280 | 3/3 |
| *T. arcticus* | Unnamed creek, NPRA | 7/11/15 | 70.235 | -151.273 | 3/3 |
| *T. arcticus* | Unnamed creek, NPRA | 7/14/15 | 70.251 | -151.778 | 3/3 |
| *T. arcticus* | Unnamed creek, NPRA | 7/14/15 | 70.255 | -151.773 | 3/3 |
| *T. arcticus* | Usuktuk river, NPRA | 7/18/15 | 70.052 | -156.548 | 3/3 |
| *T. arcticus* | Unnamed creek, NPRA | 7/20/15 | 70.513 | -155.191 | 3/3 |
| *T. arcticus* | Okpiksak River, NPRA | 7/18/15 | 70.509 | -156.455 | 3/3 |
| *T. arcticus* | Unnamed creek, NPRA | 7/20/15 | 70.515 | -155.643 | 3/3 |
| *T. arcticus* | Unnamed creek, NPRA | 7/25/15 | 70.425 | -153.326 | 3/3 |
| *T. arcticus* | Unnamed creek, NPRA | 7/19/15 | 69.859 | -160.205 | 3/3 |
| *T. arcticus* | Kucheak Creek NPRA | 7/17/15 | 70.509 | -158.066 | 3/3 |
| *T. arcticus* | Unnamed creek, NPRA | 7/14/15 | 70.620 | -157.735 | 3/3 |
| *T. arcticus* | Unnamed creek, NPRA | 7/24/15 | 70.553 | -154.518 | 2/3 |
| *T. arcticus* | Unnamed creek, NPRA | 7/25/15 | 70.276 | -154.386 | 3/3 |
| *T. arcticus* | Unnamed creek, NPRA | 7/17/15 | 70.258 | -159.613 | 3/3 |
| *T. arcticus* | Pikroka Creek NPRA | 7/15/15 | 70.129 | -157.571 | 3/3 |
| *T. arcticus* | Kukak Creek NPRA | 7/15/15 | 70.290 | -157.771 | 2/3 |
| *S. alpinus* | Lake E5, TFS | 11/5/16 | 68.642 | -149.458 | 3/3 |
| *S. alpinus* | Lake Fog1, TFS | 7/14/16 | 68.684 | -149.082 | 3/3 |
| *S. alpinus* | Lake Fog2, TFS | 11/6/16 | 68.679 | -149.091 | 2/3 |
| *S. alpinus* | Lake Fog3, TFS | 7/15/16 | 68.673 | -149.088 | 3/3 |
| *S. alpinus* | Lake Fog5, TFS | 7/15/16 | 68.678 | -149.065 | 3/3 |
| *C. cognatus* | Lake E5, TFS | 11/5/16 | 68.642 | -149.458 | 3/3 |
| *C. cognatus* | Lake Fog1, TFS | 7/14/16 | 68.684 | -149.082 | 2/3 |
| *C. cognatus* | Lake Fog2, TFS | 11/6/16 | 68.679 | -149.091 | 3/3 |
| *C. cognatus* | Lake Fog3, TFS | 7/15/16 | 68.673 | -149.088 | 1/3 |
| *C. cognatus* | Lake Fog5, TFS | 7/15/16 | 68.678 | -149.065 | 3/3 |

Water was pumped through a 25 mm, 10 μm nylon net filter (EDM Millipore; Billerica, Massachusetts; catalogue # NY1002500) using a peristaltic pump (Geotech Geopump; Denver, Colorado) until filters clogged (between 4-20 liters). Periodically during sample collection, eight total negative control field blanks consisting of 1 L of DD H_2_O carried into the field were filtered through the same equipment to test for contamination. eDNA was extracted from whole filters with a Qiagen Blood and tissue kit (Qiagen, Valencia, CA). Filters were incubated in 260 μL buffer ATL and 40 μL proteinase K for one hour at 56 °C, with vortexing every 15 minutes. Next, 300 μL buffer AT was added, followed by 300 μL 99% ethanol. Extractions then proceeded following the manufacturers recommendations, with a final elution volume of 200 μL. qPCR reactions were conducted as above, with 4 μL of eDNA extraction product as template. All rounds of extraction included at least one lab blank extraction negative control consisting of a clean filter to test for cross-contamination. All qPCRs were run in triplicate, and a minimum of six no-template negative controls were included in each qPCR run to test for contamination.

## Results

### Assay specificity and sensitivity testing

All target samples amplified in SYBR^®^-Green qPCR with their corresponding primer set. All non-target samples either did not amplify, or amplified >11 C_t_ later than targets, a range which is suitable for specificity once a selective probe is added to the assay. The melting curve produced a single sharp peak in all instances indicating no primer-dimers or off target amplification.

All target samples ampified in Taqman^®^ qPCR. Late amplification was observed for two non-target samples for the *L. lota* assays (UAM:10523 and UAM:10516), as well as four non-target samples for the *S. alpinus/S. malma* assay (UAM:6603, UAM:9910, UAM:9908 and UAM:9962) indicating either cross-contamination or cross-amplification. We aligned the primers and probe to the original sequence data generated from these samples, and all had many mismatches in both the primers and the probe, including near the 3’ end of primers, and thus cross-amplification would be highly unlikely. Upon closer examination of the UAM database, we discovered that these fin-clip samples were collected directly after a target species sample, and thus cross-contamination during fin-clip sample collection was highly likely. We sequenced qPCR amplicons from all of these cross-contaminated reactions, and all sequences matched the target species (*L. lota* or *S. alpinus/malma*) and not the species from which the actual fin-clip was taken, confirming with certainty that these samples were cross-contaminated. This affirms that care should be taken when using museum specimens for eDNA assay validation (Rodgers 2017). We ran alternate samples for each of these same non-target species that were not collected at the same time as target species, and no amplification was observed. No amplification was observed in any other non-target samples.

For all assays, all replicates amplified in qPCR down to 5 plasmid DNA copies per reaction. Thus, we are confident that these assays are highly sensitive for detection of the target species. Amplification efficiency values from the standard curve for each assay were: *L. lota* = 0.992, *C. cognatus* = 0.910, *S. alpinus/malma* = 0.999, and *T. arcticus* = 1.001.

From the NCBI Primer-BLAST Genbank searches, for the *L. lota* primer set, zero species were identified that could potentially cross-amplify. This is noteworthy, because in locations further south, such as in the Colorado River Basin, *L. lota* is an invasive species (Gardunio et al. 2011). Thus, this assay should be useful for detecting *L. lota* not only in Alaska, but also in its non-native range. For the other three primer sets, many species (mostly congeners) were identified that could potentially cross-amplify, but none that occupy the North Slope of Alaska. A full list is provided in supplementary Table S2. For the *S. alpinus/malma* primer set, numerous congeneric *Salvelinus* species native to Russia were a perfect match. Thus, our assay may be useful for detecting, but not distinguishing, Russian *Salvelinus* species as well.

### Field Validation

Amplification was observed in at least one qPCR replicate from all locations where the target species was known to be present, with the respective assay (Table 2). No amplification was observed from any of the samples tested from outside of the species range. To further confirm the identities of positive amplifications, we also sequenced eDNA qPCR products from two locations for each species. In all cases the resulting sequence matched the target species.

## Discussion

We developed 4 qPCR assays capable of detecting 5 different fish species present on the North Slope of Alaska. In testing with tissue DNA, all assays were species-specific within the North Slope fish community, with the exception of the Sal/Sma-cytb assay, which amplifies 2 closely related species, while excluding all others. In testing with known copy number plasmid DNA, all assays were highly sensitive. All assays also performed in the field for species detection from filtered water samples.

Although our Sal/Sma-cytb assay cannot distinguish between *S. alpinus* and *S. malma*, it should still be practical for most field applications due to the ecology of these two species. *S. alpinus* is almost exclusively found in lakes while *S. malma* is almost exclusively found in streams and rivers. Thus the location of detection should discriminate species in most cases, except where *S. alpinus* eDNA could potentially flow from a lake into an outlet stream. Studies of brook trout *Salvelinus fontinalis* (Jane et al. 2015; Wilcox et al. 2016) found total eDNA transport distances of <1000m from caged fish in headwater streams, although downstream transport distances are likely to be dependent on many environmental factors such as flow level, stream morphology, and eDNA starting concentration.

All of our assays were designed for use on the North Slope of Alaska; however they are likely to work in other arctic locations, other locations in Alaska and Canada, and may potentially work throughout the Arctic. In addition the *L. lota* assay should work in locations where the species is invasive in the lower 48 United States, such as the Colorado river basin (Gardunio et al. 2011). However, further specificity testing should be conducted if these assays are to be used in locations outside of the North Slope. As the majority of reference samples used to design and test our assays were from the Alaskan North Slope, it is possible that different haplotypes from other locations could produce false negatives or false positives. Additionally, if research is to be undertaken in locations where species not tested in this study are present, these additional species should be empirically tested for cross-amplification. For example, for the *C. cognatus* assay we designed primers that should, based on primer mismatches, not amplify two other *Cottus* species (*C. aleuticus* and *C. asper)* present in other regions of Alaska. However, empirical testing for cross-amplification of these two species should be conducted if studies using our assay are carried out in locations where either of these congeneric species may be present to ensure that they will not cause false positives.

Carim et al. (2016) designed an additional eDNA assay for detection of *Thymallus arcticus.* Their assay however, was designed for use in the upper Missouri river basin in Montana. Because the sympatric fish communities are quite different between the upper Missouri basin and the North Slope of Alaska (the far southern and far northern portions of the species range respectively), both assays are unlikely to be species specific in both communities. Having multiple assays for the same species, however, is beneficial as it provides managers with marker choices depending on what portion of the species range they wish to utilize eDNA for species detection.

In summary, we provide a valuable eDNA tool for detection of five Alaskan fish species that are important for subsistence fisheries in the North Slope. As North Slope populations of these species are under threat from climate change and petroleum extraction, improving knowledge of their current distributions in a timely manner is vital. eDNA methods will improve detection probability, especially for rare or low abundance species or populations. Additionally, this molecular tool will help managers obtain these important data more efficiently and economically than currently available methods, which will allow rapid and informed management decisions in the face of impending threats.

## Acknowledgements

We would like to thank Greta Burkart and the USFWS Arctic National Wildlife Refuge and Alaska Refuges Inventory and Monitoring Program for management direction and funding. We would also like to thank Andres Lopes and the University of Alaska Museum for providing tissue specimens for sequencing, and Matthew Whitman and Phaedra Budy for providing tissue specimens and eDNA samples. This work was funded by NASA Biodiversity and Ecological Forecasting program Grant NNX14AC40G.

